# Contextual and Combinatorial Structure in Sperm Whale Vocalisations

**DOI:** 10.1101/2023.12.06.570484

**Authors:** Pratyusha Sharma, Shane Gero, Roger Payne, David F. Gruber, Daniela Rus, Antonio Torralba, Jacob Andreas

## Abstract

Sperm whales *(Physeter macrocephalus)* are long-lived and highly social mammals that engage in complex group behaviours, including navigation, foraging, and child-rearing. During these behaviours, sperm whales communicate primarily using sequences of short bursts of clicks with varying inter-click intervals, known as codas. Past research has identified around 150 discrete coda types globally, with 21 in the Caribbean. A subset of these have been shown to encode information about caller and clan identity. However, almost everything else about the sperm whale communication system, including basic questions about its structure and information-carrying capacity, remains unknown. In this study, we show that codas exhibit *contextual* and *combinatorial* structure with key similarities to aspects of human language and other primate communication systems. First, we report previously undescribed variations in coda structure that are sensitive to the conversational context in which they occur. We call these *rubato* and *ornamentation*, by analogy to musical terminology. These variations are systematically controlled and imitated across individual whales. Second, we show that coda types are not defined by arbitrary sequences of inter-click intervals, but instead form a combinatorial coding system in which rubato and ornamentation combine with two categorical, context-independent features that we call *rhythm* and *tempo* to give rise to a large inventory of distinguishable codas. In a dataset of 8,719 codas from the sperm whales of the Eastern Caribbean clan, this ‘sperm whale phonetic alphabet’ makes it possible to systematically explain observed variability in coda structure. Sperm whale vocalisations are more expressive and structured than previously believed, and are built from a repertoire comprising nearly an order of magnitude more distinguishable codas. These results show context-sensitive and combinatorial vocalisation systems extend beyond humans, and can appear in an organism with a divergent evolutionary lineage and vocal apparatus.

## 1 Introduction

The social complexity hypothesis (*1, 2*) posits that animals in complex societies—featuring coordination, distributed decision-making, social recognition, and social learning of cultural strategies (*3–6*)—require complex communication systems to mediate these behaviours and interactions (*1, 7*). In humans, communication plays a particularly large role in complex social behaviours like strategising and teaching (*8–10*). These behaviours require the ability to generate and understand a vast space of possible messages. For example, the sentence *‘Let’s meet next to the statue of Claude Shannon on the fifth floor at 3pm’* picks out a specific action at a specific place and time from an enormous space of possible activities. This ability to generate complex messages, in turn, is supported by the fact that all known human languages exhibit contextual and combinatorial structure. To enable efficient communication, the meaning of most human utterances is underspecified and derived in part from the conversation that precedes them (*11*). To enable many distinct meanings to be communicated, humans access a large inventory of basic sounds (phonemes) by combining phonetic features like place of articulation, manner of articulation, then sequence phonemes to produce an unbounded set of distinct utterances (*12*–*16*). Contextuality and combinatoriality, especially below the sequence level, have few analogues in communication systems outside of human language (*17–21*). Understanding when and how aspects of human-like communication arise in nature offers one path toward understanding the basis of intelligence in other lifeforms.

Cetaceans are an important group for the study of evolution and development of sophisticated communication systems (*22*). Among cetaceans, long-term observational studies of sperm whales *(Physeter macrocephalus)* have described both a culturally defined, multi-level, matrilineal society (*23*) and a socially transmitted communication system (*24–26*). Sperm whales are known for complex social and foraging behavior, as well as group decision-making (*27*). They communicate using **codas** (*28*): stereotyped sequences of 3-40 broadband clicks. Codas are exchanged between whales when socialising or between long, deep, foraging dives (*23*). To date, scientists have described the sperm whale communication system in terms of a finite repertoire of **coda types**, each defined by a characteristic sequence of **inter-click intervals** (ICIs) as seen in Fig. 1(A). Coda type repertoires can be defined manually or using automated clustering procedures, and have been used to delineate cultural boundaries among socially segregated but sympatric ‘clans’ of whales (*24*) whose members differ in their behavior (*25,29,30*). But there is an apparent contradiction between the social and behavioral complexity evinced by sperm whales and the comparative simplicity of a communication system with a small, fixed set of messages. This contradiction naturally raises the question of whether any additional, previously undescribed structure is present in sperm whale vocalisations.

**Figure 1:**
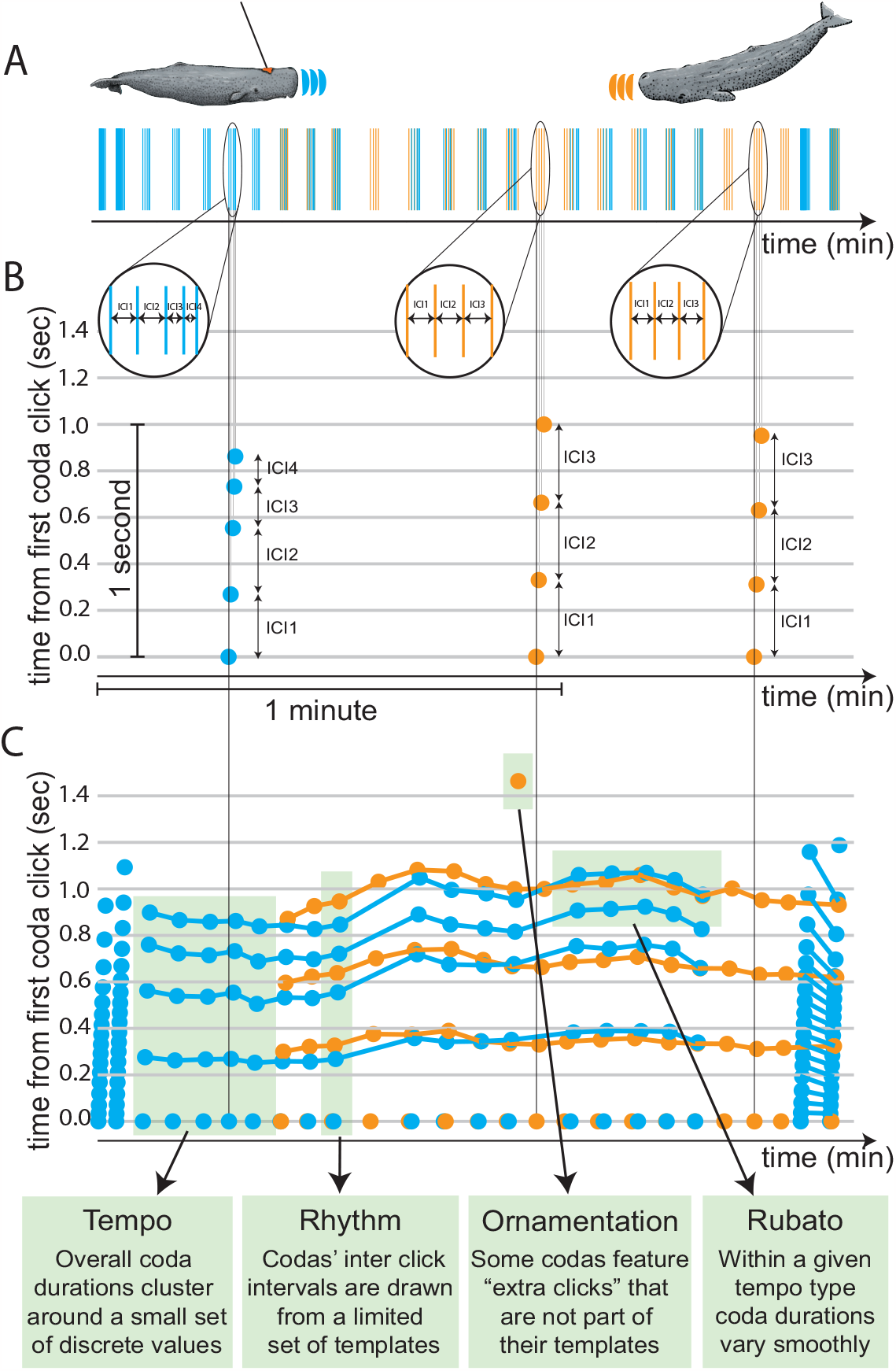
Exchange Plot: Sperm whales communicate by producing sequences of clicks. **(A)** shows a two-minute exhange between two whales (with clicks visualized in blue and orange respectively) from the Dominica Sperm Whale Dataset. **(B)** projects these clicks over a time– time plot, in which the horizontal axis plots the time in the exchange at which a click occurs, and the vertical axis represents the time of the click from the first click in the coda. The vertical axis serves as a microscope over the horizontal axis, revealing the internal structure of each coda. **(C)** shows a time–time visualization for the entire two-minute exchange (with lines connecting matching clicks between adjacent codas), revealing complex, context-dependent variation in coda structure.

We first demonstrate that some coda structure is contextual:^1^ when analysing codas exchanged between whales, we observe fine-grained modulation of inter-click intervals relative to preceding codas, as well as modification of standard coda types via the addition of an extra click. We term these contextual features **rubato** and **ornamentation**. Next, we show that the coda repertoire has combinatorial structure: in addition to rubato and ornamentation, codas’ **rhythms** and **tempos** can independently be discretised into a small number of categories or types. We show that all four features are sensed and acted upon by participants in the vocal exchanges, and thus constitute deliberate components of the whale communication system rather than unconscious variation. Rhythm, tempo, rubato and ornamentation can be freely combined, together enabling whales to systematically synthesize an enormous repertoire of distinguishable codas. While the communicative function of many codas remains an open question, our results show that the sperm whale communication system is, in principle, capable of representing a large space of possible meanings, using similar mechanisms to those employed by human sound production and representation systems like speech, Morse code, and musical notation.

## 2 Analysis

### The Dataset

For this study, we used the annotated coda dataset from The Dominica Sperm Whale Project (DSWP), the current largest sperm whale data repository. The recordings of the Eastern Caribbean 1 (EC-1) clan were used in the analysis, comprising 8,719 codas making up 21 previously defined coda types. This dataset contains manually annotated coda clicks and extracted inter-click intervals in data recorded from various platforms and various recording systems between 2005 and 2018, including animal-worn, acoustic, biologging tags (D-tags) deployed between 2014 and 2018 (see Supplementary 2). The EC-1 clan comprises 400 individuals. 42 tags were deployed on 25 different individuals in 11 different social units. Three of these are less-studied units, for which precise size estimates are not available. We conservatively estimate that at least 60 distinct whales are recorded in the DSWP dataset.

### Exchange Plots: Visualizing multi-whale calls

Codas, considered to be the basic units of sperm whale communication, consist of click groups generally less than two seconds in duration. Previous work has defined coda types and characterised coda repertoires by examining single codas outside the context in which they were produced. However, codas are not produced in isolation, but in interactive exchanges between two or more chorusing whales. Individual whales within a chorus tend to produce sequences of codas with a periodicity of approximately 4 seconds (see Supplementary Section 3). Across a chorus, interacting whales produce codas both alternately (i.e. turn-taking) or near-simultaneously (i.e. overlapping). Therefore, sperm whale vocalisations demonstrate complexity on two different timescales: a fine-grained timescale that determines the makeup of each individual coda, and a longer time scale that determines the overall structure of the interactive exchange across codas within a chorus.

We depict these vocalisations using a new visualisation we call an **exchange plot** (Fig. 1 (B-C)). Both axes of this plot measure time: the horizontal axis shows the time elapsed since the beginning of a conversation, and the vertical axis shows the time since the onset of each coda. Exchange plots reveal the structure of codas in their interactive context, making it possible to observe both fine-grained differences between adjacent codas made by interacting whales, and long-range trends over the course of an exchange.

### 2.1 Contextuality

Visualising whale vocalisations with an exchange plot, as in Fig. 1(B-C), makes it apparent that characterising sperm whale interactions as sequences of fixed coda types overlooks a great deal of information. First, coda duration varies smoothly over the course of an exchange; variation in the duration of a whale’s codas is systematically imitated by interlocutors, even when coda-internal click spacing differs (Fig. 1(C)). Second, some ‘extra’ clicks in Fig. 1(C) appear at the end of codas that otherwise match their neighbors. We hypothesised that these smooth variations and extra clicks constitute perceptible and controllable features of codas independent of their discrete type, pointing toward a more complex sperm whale communication system with a greater information-carrying capacity than previously reported.

#### Rubato

**Codas exhibit fine-grained duration variation in addition to discrete tempo** A coda’s **duration** is the sum of its inter-click intervals Fig. 2(A). While durations tend to cluster around a finite set of values Fig. 2(A), there remains substantial continuous variation within these clusters. Past work has described these differences as unexplained ‘within-type variation’ of categorical coda types (*24, 32*). However, the *structured* nature of this variation—which, as shown in Fig. 1(C), is temporally correlated and imitated across whales—has never been previously documented. We demonstrate that variation in coda duration is not random: individual whales modulate coda durations smoothly over the course of multi-coda exchanges without necessarily imitating click counts or inter-click interval spacing. An example is depicted in Fig. 1(C): one whale produces a sequence of codas smoothly varying in duration, while its interlocutor closely matches these changes in overall coda duration but not the number of clicks.^2^

**Figure 2:**
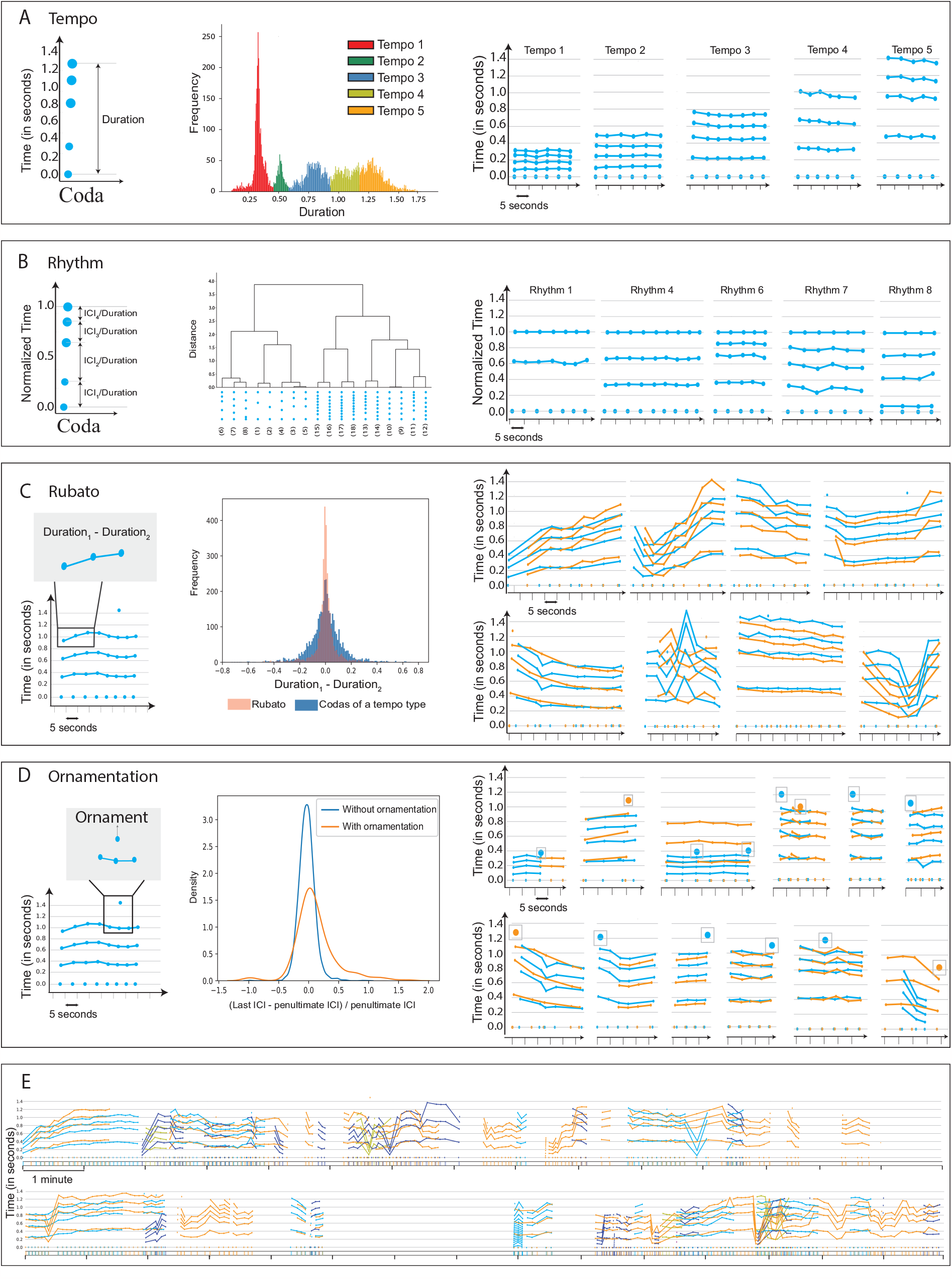
Combinatorial basis of coda production: Sperm whale codas were previously hypothesized to comprise 21 discrete types. We show that this coda repertoire is instead built from two context-independent features (rhythm and tempo) and two context-sensitive features (rubato and ornamentation). **(A) Tempo:** (*Left*) The overall duration of a coda is the sum of its interclick intervals. (*Centre*) Coda durations are distributed around a finite set of modes, which we call *tempo types*. (*Right*) Snippets of exchanges showing different tempo types. **(B) Rhythm:** (*Left*) Normalizing the vector of ICIs by the total duration returns a duration-independent coda representation, which we call rhythm. (*Centre*) Codas cluster around 18 rhythm types. (*Right*) Normalized codas showing different rhythm types. **(C) Rubato:** (*Left*) Sperm whales slowly modulate coda duration across consecutive codas, a phenomenon we call *rubato*. (*Centre*) Rubato is gradual: adjacent codas have durations more similar to each other than codas of the same type from elsewhere in an exchange. (*Right*) Whale choruses with imitation of rubato. **(D) Ornamentation:** (*Left*) Some codas feature ‘extra clicks’ (*ornaments*) not present in neighboring codas that otherwise share the same ICIs. (*Centre*) A density plot showing the distribution of the ratio between final ICIs in ornamented codas versus un-ornamented codas. Ornamented codas have a significantly different ICI distribution compared to regular codas. (*Right*) Examples of ornaments. **(E) Thirty minutes of multi-whale choruses:** Exchanges feature imitation of coda duration across whales, gradual changes in call structure, and rich contextual variability.

By analogy to the corresponding musical phenomenon, we call this behavior **rubato**.

To show that rubato is not random variation, we first evaluated whether changes in duration are smooth by measuring whether codas are more similar to their neighbors than other codas of the same type. To do so, we computed the **tempo drift** between two codas from the same speaker, defined as the difference in coda durations (Fig. 2(C)). We computed the average absolute drift between (1) adjacent coda pairs and (2) random coda pairs of the same discrete coda type (according to its rhythm and tempo; see Section 2.2 for a discussion of rhythm and tempo and Supplementary Section 6 for additional details). We found that drift was significantly smaller between adjacent codas (0.050s on average) than would be expected under a null hypothesis that drift depends only on a coda’s discrete type (which would give a drift of 0.100s on average; test: permutation test (one-sided), *p* = 0.0001, *n* = 2593; see Supplementary Material for details). Thus, within-type variation is context-dependent.

Second, we evaluated whether sequences of codas reflect longer-term trends. To do so, we collected coda triples of the same discrete coda type, and measured the correlation between tempo drift across adjacent pairs. We found a significant positive correlation, compared to a null hypothesis that drift between adjacent pairs is uncorrelated (test: Spearman’s rank-order correlation (two-sided), *r*(2586) = 0.57, *p* = 2*e*^*−*220^, 95% CI= [0.54, 0.60], *n* = 2588). Thus, rubato is distributed across sequences of multiple codas.

Finally, we evaluated whether rubato is perceived and controlled by measuring whales’ ability to match their interlocutors’ coda durations when chorusing. We measured the average absolute difference in duration between (1) pairs of overlapping codas from different whales, and (2) pairs of non-overlapping codas of the same discrete coda type. Durations are significantly more closely matched for overlapping codas (0.099s on average) than would be expected under a null hypothesis that chorusing whales match only discrete coda type (which would give a drift of 0.129s on average) (test: permutation test (one-sided), *p* = 0.0001, *n* = 908; see Supplementary Section 6).

#### Ornamentation

**Some clicks do not belong to standard tempo types** Fig. 2(D) depicts an exchange consisting of one six-click coda, followed by a long sequence of five-click codas. The first five clicks of the initial 6-click coda closely match those of neighboring codas: if the sixth click were removed, we would identify the first coda as having nearly the same inter-click intervals as its neighbors (and assign it to the same discrete coda type). While not previously described, ‘extra clicks’ of this kind occur in (4.6%) of the codas in the exchanges of Eastern Caribbean whales. Additional examples appear in Fig. 2(D) and Supplementary Section 7. We hypothesised that ‘extra’ clicks play a different role from the other clicks in the codas in which they appear: they do not determine discrete coda type. Instead, like rubato, they constitute an independent feature of the sperm whale vocalisation system. We call these extra clicks **ornaments**.

We define an ornament as the final click in a coda containing one more click than the nearest preceding or following coda within a window of ten seconds, with this interval being based on the average response time (Supplementary Section 3). To show that these ornaments play a distinct role from other clicks, we first computed the mean squared distance between each standardised, ornamented coda and the cluster centre of its assigned rhythm type. We then removed ornaments and computed mean squared distance between the standardised coda and the centres of rhythm clusters for adjacent codas produced by the same whale. The second quantity (0.0034s^2^, reflecting a hypothesis that ornamented codas match their neighbors) is significantly smaller than the first (0.0053s^2^, reflecting a null hypothesis that ornamented codas resemble other codas of the same type) (test: permutation test (one-sided), *p* = 0.002, *n* = 178). In other words, ornamented codas are *less* like other codas with a matching number of clicks, and *more* like neighboring non-ornamented codas, if ornaments are modeled as a separate factor. To further verify that ornaments are distinct from other clicks, we compared ICIs (inter-click intervals) in ornamented vs. non-ornamented codas with the same number of clicks. We specifically compared the difference between the final two ICIs, normalized by the penultimate ICI, to reduce variance arising from rubato. This measurement exhibits a significant difference in distribution in ornamented vs non-ornamented codas (test: Kolmogorov-Smirnov test (two-sample), *D*(178, 3666) = 0.28, *p* = 2*e*^*−*14^, 95% CI = [0.17, 0.39], *n* = (178, 3666), Fig. 2(D)).

Finally, ornaments are not distributed randomly, but appear in distinctive positions in longer exchanges. Within a single whale’s call sequences (defined as a sequence of codas separated by no more than eight seconds), a significantly greater proportion of ornamented codas appear at the beginning of call sequences than unornamented codas (test: Fisher’s exact test (two-sided), odds ratio: 2.00, *p* = 0.0006). A significantly greater fraction of ornamented codas also appear at the *end* of call sequences compared to unornamented codas (test: Fisher’s exact test (twosided), odds ratio: 2.07, *p* = 0.0004). Moreover, ornaments are are predictive of changes in multi-whale interactions. We define a ‘change in chorusing behavior’ as one of three events: a following whale begins chorusing with a leading whale, pauses chorusing, or ceases vocalizing for the remainder of the exchange. Compared to unornamented codas, ornamented codas from the *leading* whale are disproportionately succeeded by a change in chorusing behavior from the *following* whale (test: Fisher’s exact test (two-sided), odds ratio: 1.91, *p* = 0.0007). Thus ornamentation, like rubato, appears to be perceived in multi-whale interactions.

### 2.2 Combinatoriality

The existence of ornamentation and rubato features demonstrates that codas carry more information, and have more complex internal structure, than their discrete type alone would indicate. Instead, they result from a combinatorial coding system in which discrete type, ornamentation and rubato combine to realise individual codas. Based on these findings, we hypothesized that categorical coda types might themselves arise from a combinatorial process, with a simpler set of features explaining the prototypical ICI vector for each coda type.

During rubato, consecutive calls from a single whale gradually vary coda duration while preserving the relative relationship of ICIs (Supplementary Section 4.3), suggesting that whales are capable of maintaining this relationship independent of its duration. Moreover, existing studies have noted that codas may also be discretely clustered according to standardised ICI (*24,32,34*), a process that assigns codas with very different durations to the same cluster. Finally, some instances of chorusing involve whales producing codas with different ICIs (or different numbers of clicks) but matched durations. Together, these observations suggest that past inventories of discrete coda types (e.g. (*35*)) might specifically be interpretable in terms of two features: a normalised ICI category (which we term a coda’s **rhythm**) and a discrete duration category (independent of rubato, and which we term **tempo**). To validate this hypothesis, we measured (1) whether coda rhythms and tempos cluster around a discrete set of values, and (2) whether rhythm and tempo features are independently combinable (both with each other and with ornamentation and rubato features).

As shown in Fig. 2(A), codas with the same duration may have different internal click spacing (and even different numbers of clicks) but still span the same amount of time from the first click to the last click. Performing kernel density estimation (KDE) on scalar coda durations from the DSWP dataset reveals five distinct modes in the distribution of durations (Fig. 2(A)), indicating that the number of realized coda durations is much smaller than the total number of identified coda types (Supplementary Section 5).

Across codas, the relative relationships between ICIs are often repeated even independent of tempo. In Fig. 2(A), for example, we see two five-click codas, one long and one short, but both characterised by the uniform spacing of the constituent ICIs. Past work has shown that these **rhythms** are reused; our analysis uses the 18 rhythm clusters proposed by (*35*) (detailed breakdowns are given in Fig. 2(B) and in Supplementary Section 4).

Finally, to evaluate the combinability of these features, we computed the frequency with which each rhythm and tempo feature co-occurred in the DWSP dataset, as well as the frequency with which each combination appeared with ornamentation or rubato. Results are shown in Fig. 3. Each rhythm type appears with at least one tempo types and each tempo type appears with at least three rhythm types. Moreover, (22%) of these combinations can appear with or without rubato and ornamentation.

**Figure 3:**
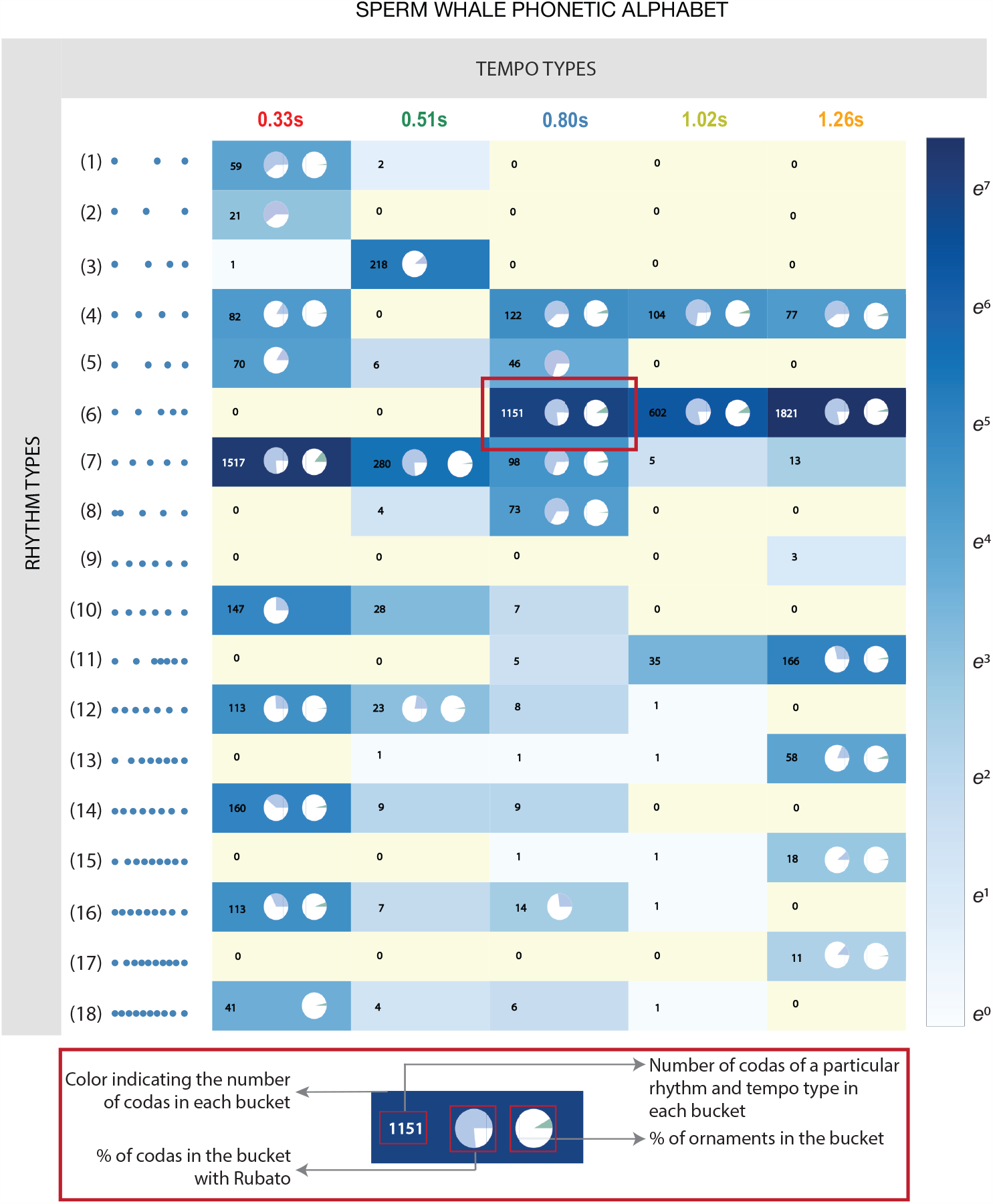
Sperm Whale Phonetic Alphabet: Analogous to visualizations of the human phonetic repertoire, we propose a phonetic alphabet for sperm whales. Tempo types are plotted on the vertical axis, rhythm types are plotted on the horizontal axis, and the color of each cell represents the number of occurrences of that rhythm/tempo combination in the DSWP dataset. Pie charts in each cell provide further information about the prevalence of rubato and ornamentation within each combination: the left pie shows the ratio of the number of codas that appear with rubato to those without, while the right pie shows the fraction of all ornaments that appear with that feature combination. While not all feature combinations are realised (as observed in human languages), sperm whale codas have a rich combinatorial structure with both discrete and continuous parameters and at least 143 combinations frequently observed (Supplementary Section 8).

Like the International Phonetic Alphabet for human languages, this ‘Sperm Whale Phonetic Alphabet’ (Fig. 3) shows how a small set of axes of variation (place of articulation, manner of articulation, and voicedness in humans; rhythm, tempo, ornamentation, and rubato in sperm whales) give rise to the diverse set of observed phonemes (in humans) or codas (in sperm whales). As in human languages, not all theoretically realisable feature combinations are attested in the DSWP dataset, and some combinations are more frequent than others. As in human languages, most coda variation is discrete: though ICIs can vary continuously in principle, only specific patterns (associated with specific rhythms and tempos) are realised in practice. Supplementary Section 4 shows the full set of codas in the dataset, organised by rhythm, tempo, and the presence of rubato and ornamentation for each combination of rhythm and tempo. Notably, these factors of sub-coda variation exist alongside another combinatorial process—the sequential ordering of codas shown in Figure 1(E)—in which codas of different types are combined in sequence to give rise to an even larger family of distinct vocalisations, reminiscent of the bi-level combinatorial structure of speech production in humans.

Fig. 3 also demonstrates that these vocalisations have a significantly greater information capacity than was previously known. Prior work identified 21 discrete coda types, and the system could be understood to have an information rate of at most 5 bits/coda. However, our analysis suggests that with 18 rhythms, 5 tempos, optional ornamentation, and three variations (increasing, decreasing or constant duration) in rubato, the information rate could be up to twice as large (details in Supplementary Section 8). The role of rubato within this coding system remains an important open question: it might be discrete (with some simpler inventory of contours explaining the patterns in Fig. 1(C) and Fig. 2(C), as in the songs of birds (*36–41*), and humpback whales *(Megaptera novaeangliae)* (*42, 43*)). Or it might convey continuous-valued information, analogous to the orientation and duration features of the waggle dance in bees *(Apis sp*.) (*44*).

## 3 Discussion

Our results demonstrate that sperm whale vocalisations form a complex combinatorial communication system: the seemingly arbitrary inventory of coda types can be explained by combinations of rhythm, tempo, rubato, and ornamentation features. Sizable combinatorial vocalisation systems are exceedingly rare in nature; however, their use by sperm whales shows that they are not uniquely human, and can arise from dramatically different physiological, ecological, and social pressures.

These findings also offer steps towards understanding how sperm whales transmit meaning. In some organisms with combinatorial codes, such as honey bees *(Apis sp*.), the constituent features of the code transparently encode semantics (e.g. direction and distance to food sources). Further research on sperm whale vocalisations may investigate if rhythm, tempo, ornamentation, and rubato function similarly, directly encoding whales’ communicative intents. Alternatively, one of the key differentiators between human communication and all known animal communication systems is *duality of patterning:* a base set of individually meaningless elements that are sequenced to generate a very large space of meanings. The existence of a combinatorial coding system—at either the level of sounds, sound sequences, or both—is a prerequisite for duality of patterning. Our findings open up the possibility that sperm whale communication might provide our first example of that phenomenon in another species.

## Acknowledgements

We thank the Chief Fisheries Officers and the Dominica Fisheries Division officers for research permits and their collaboration in data collection; all the crews of R/V Balaena and The DSWP team for data collection, curation, and annotation; as well as Dive Dominica, Al Dive, and W.E.T. Dominica for logistical support while in Dominica.

## Author Contributions

PS JA AT DR SG RP conceptualized the study, PS JA AT developed the methods, SG and The DSWP team collected the data, SG and The DSWP team annotated the original dataset, PS SG and The DSWP team curated the data, PS conducted the analyses and authored the code provided, PS JA AT SG verified the analytical results, SG DG DR AT JA funded the study, PS wrote the manuscript, all authors contributed review and editing to the manuscript and gave final approval for publication.

## Data Availability

Data from The Dominica Sperm Whale Project were collected under scientific research permits from the Fisheries Division of the Government of Dominica. The field protocols for approaching, photographing, tagging, and recording sperm whales were approved by either the University Committee on Laboratory Animals of Dalhousie University, Canada; the Animal Welfare and Ethics Committee of the University of St Andrews, Scotland; or Aarhus University, Denmark; and sometimes several or all of these across years. The dataset and custom scripts used for this study will be made available upon publication.

## Footnotes

1 Throughout this paper, we use *context* to denote conversational context (e.g. neighboring codas) rather than behavioral context (e.g. diving), as is standard in the study of natural and formal languages (*31*).

2 This refines the finding in (*33*) that overlapping codas were more likely to be of the same coda type than expected by chance: sometimes codas share a duration but not a discrete type.

