## Supplement for "Contextual and Combinatorial Structure in Sperm Whale Vocalisations"

### Supplementary Information: Contextual and Combinatorial Structure in Sperm Whale Vocalisations

September 2023

#### 1 Glossary

1. **Coda:** *A short burst of clicks with varying inter-click intervals generally less than two seconds in duration.*

2. **Inter Click Interval (ICI):** *The time difference between two consecutive clicks within a coda.*

3. **Absolute ICI:** *The absolute time difference between consecutive clicks in a given a coda as produced by the whale and recorded (see (1)).*

4. **Coda duration:** *The sum of a coda's absolute ICIs.*

5. **Standardised absolute ICI:** *ICI normalized by the total duration of the containing coda. This conserves rhythm but discards tempo (see (1)).*

6. **Cumulative ICI:** *The absolute time difference between any given click and the first click of a coda as produced by the whale and recorded.*

7. **Standardised cumulative ICI:** *Relative ICI normalized by the total duration of the containing coda. This conserves the rhythm of the coda but discards the tempo. In codas normalized in this way, the last relative ICI is equal to 1 (compare to standardized absolute ICIs, which* sum *to 1).*

8. **Coda Type:** *Categorical coda representation (primarily used in past work) obtained by clustering codas according to absolute ICI, which accounts for both rhythm and tempo simultaneously.*

9. **Rhythm type:** *The discrete category a coda is assigned to based on its characteristic sequence of standardized ICIs.*

10. **Tempo type:** *The discrete category a coda is assigned to based on its characteristic duration.*

11. **Exchange / Chorus:** *Period of time where codas are made by more than a single whale (as in (2)).*

12. **Single-Whale Call Sequence:** *A sequence of calls made by a given whale where every consecutive pair of calls occur within 8 seconds (twice the average response time) of each other.*

13. **Turn-taking:** *An exchange of codas involving alternating coda production. Also referred to as 'adjacent' codas, these are defined as next-in-sequence codas whose onset occurred within two seconds, but after the termination, of the initial coda (as in (3)).*

14. **Overlapping Codas:** *An exchange of codas such that the next-in-sequence coda's onset occurs after the onset, but before the termination, of the previous coda (as in (3)).*

15. **Ornament:** *"Extra click" appended to the end of a coda in a group of shorter codas. (For further details on the identification criterion, see Ornamentation section in the manuscript.)*

16. **Rubato:** *Gradual variation in duration across adjacent codas made by the same whale within the same rhythm and tempo type.*

#### 2 Data Collection and Coda Annotation

Social units of female and immature sperm whales were located and followed in an area that covered approximately 2000 squared kilometers along the entire western coast of the Island of Dominica (N15.30 W61.40) between 2005 and 2018.

Codas were recorded using one of several recording setups: In 2005, we used a Fostex VF-160 multitrack recorder (44.1 kHz sampling rate) and a custom-built towed hydrophone (Benthos AQ-4 elements, frequency response: 0.1–30kHz) with a filter box with high-pass filters up to 1 kHz resulting in a recording chain with a flat frequency response across a minimum of 2–20kHz. No recordings were made during the short 2006 season. In the 2007, 2009, 2011, 2016, and 2017 seasons, we used a Zoom H4 portable field recorder (48 kHz sampling rate) and a Cetacean Research Technology C55 hydrophone (frequency response: 0.02-44kHz) and no filters. During the 2008, 2010, 2012, 2015, and 2018 seasons, we used a custom-built towed hydrophone (Benthos AQ-4 elements, frequency response: 0.1-30kHz) with a filter box with high-pass filters up to 1 kHz resulting in a recording chain with a flat frequency response across a minimum of 2-20 kHz. This was connected to a computer-based recording system as a part of the International Fund for Animal Welfare's (IFAW) LOGGER software package (48 kHz sampling rate) or PAMGUARD (minimum 48 kHz sampling rate) (*4*).

In addition, recordings were also made through the deployment of animal-borne sound and movement tags (DTag generation 3, Johnson and Tyack 2003). Tagging was undertaken between 2014 and 2018 on an 11-meter rigid-hulled inflatable boat (RHIB). Tags were deployed from a 9-meter, hand-held, carbon fiber pole, and were attached to the whales using four suction cups. DTags record two-channel audio at 120 kHz with a 16-bit resolution, providing a flat (±2 dB) frequency response between 0.4 and 45kHz. Pressure and acceleration were sampled at a rate of 500 Hz with a 16-bit resolution, and were decimated to 25 Hz for analysis. DTag analysis was conducted using custom scripts in Matlab 2015b (The Mathworks, Inc., MA, USA). The variation in the frequency responses and sampling rates of the recording systems used did not affect our ability to record clean signals for both the coda and echolocation clicks produced by sperm whales, and as a result, the temporal patterning of clicks used in this analysis.

Whales, including the tagged whales, were identified through photographs of the trailing edge of their tails (*5*). Identifications were used to ensure that only recordings from one of the two sympatric clans (EC1, the Eastern Caribbean Clan) were included in the analysis to control for any differences in repertoires between vocal clans (*6*).

To define the temporal structure of the codas recorded, absolute inter-click intervals were measured as in (*7*), using either custom-written Matlab tools and Rainbow Click Software (all years before 2014) or Coda Sorter a custom-written tool (K. Beedholm, Marine Bioacoustics Lab, Aarhus University) in LabView (National Instruments, TX, USA). CodaSorter allows users to play back audio at various speeds and manually mark detected clicks as belonging to a specific coda. Estimates for each click for the angle of arrival, channel delay, centroid frequency, and inter-pulse interval (IPI, the time between the onset of the first pulse and the onset of the next pulse in the multi-pulse structure of sperm whales clicks, (*8*)) allowed for determining if the codas were produced by tagged whales or non-focal animals; and to ensure that, on days in which multiple tags were deployed, codas recorded by different tags were not double-counted. Photo-identification supported this process by identifying which whales were present and associated with the tagged whales at each surfacing.

There are two components to the dataset: Dataset 1, a large set of all 8719 codas that are annotated with information on their inter-click intervals; and Dataset 2, a smaller set of 3948 codas, which were recorded from animal-borne DTags, which remain in temporal order and are additionally annotated with information of the absolute time in the day of the first click of each of the codas and their associated speaker identities across the bouts. Experiments that do not require contextual information (those discussing the context-independent features of rhythm and tempo) use Dataset 1, whereas those requiring information about the relative ordering of the codas and their speaker IDs (those discussing the context-sensitive features of rubato and ornamentation), use Dataset 2. In both cases, rare, long codas were excluded from analysis (greater than 10 clicks, less than 5% of all codas recorded).

### 3 Preliminaries

Consider a coda $\mathbf{c}$ consisting of five clicks occurring at times $[t_1, t_2, t_3, t_4, t_5]$. We may represent this coda as a vector of four *inter-click intervals* $c$, writing:

$$\mathbf{c} = [c_1, c_2, c_3, c_4] = [t_2 - t_1, t_3 - t_2, t_4 - t_3, t_5 - t_4] \ . \tag{1}$$

An example is depicted in Fig. 1(B) of the main paper. This coda consists of two long ICIs followed by two short ICIs. The coda may also be represented by its *cumulative inter-click intervals*:

$$\mathbf{c}_{REL} = [t_2 - t_1, t_3 - t_1, t_4 - t_1, t_5 - t_1] \ . \tag{2}$$

The exchange plot visualization plots the $i^{th}$ click of the coda such that,

$$(x, y) = (t_i, t_i - t_1) \ . \tag{3}$$

(75%) of the codas used by sperm whales of the Eastern Caribbean clan are 5-click codas. Fig. 1 shows the frequency distribution of click count.

The periodicity of codas produced by a single whale is fairly isochronous with speakers making codas with an interval of approximately four seconds. During exchanges between whales or in chorus, the inter-coda interval is bimodal, with some codas being overlapped by their interlocutor and others being produced in sequence at an interval of approximately four seconds (Fig. 2).

### 4 Rhythm

A coda's rhythm is determined by its characteristic sequence of standardized ICIs. We use the rhythm clusters shown in Fig. 3, reported by (*9*). While (*9*) has provided intrinsic (within-coda) evidence that rhythm types are clustered, this clustering is also reflected at the exchange level, between adjacent codas. We present two pieces of temporal evidence that indicate the need for an independent treatment of the overall duration of the coda from its underlying rhythm.

##### 4.1 Transition dynamics of consecutive calls

The plot in Fig. 4 shows the probability of transitioning between codas of various rhythm types by a single whale. Transitions between adjacent codas of the same rhythm type are significantly more likely than cross-type transitions, making up (82%) of all transitions. This is also clear from exchange plots, in Fig. 1 of the main paper, in which a given whale can be seen to maintain its rhythm in the course of producing consecutive calls. A high probability of transitioning into the same rhythm cluster in consecutive turns indicates that discovered clusters are also coherent in time.

##### 4.2 Overlapping codas with distinct rhythm types still match overall durations

Sometimes, overlapping codas exhibit different rhythm types (as shown in Fig. 5) and even different numbers of clicks. For example, (14%) of overlapped codas involve one 4-click coda and one 5-click coda. Even when rhythm type is not matched, duration often is matched over the course of an exchange: the difference between overlapping coda durations is on average 0.104 seconds. Under a null hypothesis that this duration matching is fully explained by the overlapping codas' discrete types, we would expect a difference in durations of 0.134 seconds, which is significantly larger(test: permutation test (one-sided), $p = 0.0001$, $n = 248$, performed by randomly resampling durations within each discrete coda type without replacement). Thus, rhythm and tempo features can be independently combined:[1] tempo can be imitated even while holding different rhythm types constant.

#### 5 Tempo

A coda's tempo is a discretised version of its overall duration.

##### 5.1 Clustering

We obtained the empirical distribution of coda durations from the DSWP dataset, then estimated the probability density function of the duration using a kernel density estimator (KDE) with a Gaussian kernel function. The bandwidth of the estimator (h=0.035) was chosen such that it minimizes the mean integrated squared error. The empirical distributions of codas cluster around a small set of modes as seen in Fig. 2(B) of the main manuscript. Any further reduction in the bandwidth causes the estimator to produce a larger set of modes in regions with insufficient data points, such as regions with duration $(< 0.1)$ seconds and duration $(> 1.5)$ seconds.

[1] We emphasize that this does not imply that rhythm and tempo are statistically independent: indeed, codas with specific rhythm types are more likely to have certain tempo types than others.

This estimated distribution discovers a set of 5 peaks with values at $[0.33, 0.51, 0.8, 1.02, 1.26]$, and end points at $[0.45, 0.61, 0.93, 1.08]$.

Several factors justify the choice of five discrete tempo clusters rather than a smaller number. Merging tempo clusters 4 and 5 would result in a bi-modal distribution of durations *within* the final tempo cluster. Merging clusters 1 and 2 causes the distribution of rubato to be bimodal only for the (new) tempo cluster 1 as shown in Fig. 9. Thus, any system with a smaller number of tempo clusters either introduces (1) a bimodal tempo distribution within the cluster or (2) a bimodal rubato distribution only within that cluster. Each of these alternatives produces a system with a single idiosyncratic cluster, in contrast to the five-cluster system.

### 6 Rubato

The rubato at any given point in the exchange is characterized by the difference in the durations of adjacent codas of the same rhythm and tempo type made by the same whale. The experiments below provide evidence that (1) whales gradually vary the total duration of their codas in consecutive calls, (2) gradual drifts over a period cause long-term changes in overall duration, and (3) this drift in duration is perceived and imitated by whales in the exchange.

#### 6.1 Adjacent coda pairs have small duration differences

The durations of adjacent codas (of the same rhythm type made in consecution by a given whale) are similar, differing by an average of 0.050s. Under a null hypothesis that duration depends only on discrete coda type, and not adjacent codas, we would expect a significantly larger value of 0.100s (test: permutation test (one-sided), $p = 0.0001$, $n = 2953$, performed by randomly resampling durations within each discrete coda type without replacement).

Results are shown in the main paper Fig. 2(C).

#### 6.2 Rubato accumulates over time, causing larger changes in duration

Changes in coda duration are positively correlated across adjacent coda triples. We compare $H_0$: consecutive rubato changes are uncorrelated, vs. $H_1$: consecutive changes in rubato are correlated and reflect longer-term trends (test: Spearman's rank-order correlation (two-sided), $r(2586) = 0.57, p = 2e^{-220}$, 95% CI= $[0.54, 0.60]$, $n = 2588$). Results are discussed in Section 2.1 of the main paper.

#### 6.3 Durations of overlapping codas from different whales are imitated

For overlapping codas, the overall duration of the coda of the initiating whale is similar to the duration of the coda produced by the interlocutor: see Fig. 10. The mean observed difference in durations is 0.99s. Under a null hypothesis that chorusing whales only match discrete coda type, we would expect a significantly larger difference of 0.129 (test: permutation test (one-sided), $p = 0.0001$, $n = 908$, performed by randomly resampling durations within each discrete coda type without replacement).

#### 6.4 Precision of imitation

Fig. 11 (*Left*) shows the distribution of the difference in duration of overlapping codas and Fig. 11 (*Right*) shows the cumulative distribution function of the absolute difference in the duration of overlapping codas. It can be seen that (45%) of the codas' duration is imitated with a precision of less than 0.1 seconds.

When overlapping codas are made, the coda duration is determined by the leading whale, and its interlocutor has to determine the timing of its clicks before the complete coda of the leading whale can be observed. Therefore, the interlocutor has to decide when to produce the $n^{th}$ click (where $(n > 1)$) of its coda after the leading whale has produced only the first $m$ clicks of its coda (where $(m > 1)$). To do this, the interlocutor is required to estimate the duration of the leading whale's coda on the basis of just the first $m$ prefix clicks of the coda.

In (92%) of overlapped codas, the first click of the following whale is made between the first and second click of the leading whale. In (97%) of overlapped codas, the second click of the following whale is made between the second and third click of the leading whale. Estimating the duration of the leading whale's coda thus requires the second whale to both precisely measure its first ICI, and additionally infer the rhythm type of the leading whale's incompletely produced coda. Since the consecutive codas of a given whale are likely to be of the same rhythm type (see Section 4), information on the rhythm type of the leading whale is present in previously produced adjacent calls. A close matching of the overall duration by the interlocutor despite an incomplete observation of the leading whale's coda demonstrates the interlocutor's awareness of the leading whale's rhythm type.

#### 6.5 Rubato is not correlated with body motion

The same tags used to record vocalization are also instrumented with accelerometers and gyroscopes, making it possible to test for correlation between vocalization and movement. Figure 13 depicts the correlation between rubato and the orientation and acceleration of the whale recorded by sensors. We find no significant correlation. The correlation between the quantities was checked from orientation and acceleration measurements made zero seconds apart to motion readings up to 10 seconds preceding the rubato. The correlation was computed for measurements in intervals of 0.1 seconds between 0 and 10. The lack of significant correlation rules out the effect of synchronized motion being the cause of coordinated rubato change.

### 7 Ornamentation

Ornamented codas are anomalous in terms of the number of clicks, duration, as well as in the aspect of their rhythm from the surrounding set of codas. In this section, we provide extra information about experiments that highlight additional differences between ornamented codas and non-ornamented codas: (1) Discarding the ornament of a coda causes the coda constituted by the remaining clicks to match the rhythm type of its neighbouring codas. (2) The rubato of ornamented codas has a distribution different from non-ornamented codas. (3) ICIs of ornamented codas have anomalous statistics. Finally, (4) ornaments are not imitated by whales in a chorus.

#### 7.1 Finding ornaments

We define an ornament as the last click of the coda that contains one more click than the immediate neighbouring codas made by the same whale. In the dataset, we find that (4.6%) of the codas contain an ornament.

Since ornament labels are assigned purely based on the number of clicks, the effects on rhythm, and duration described below are not necessary consequences of this procedure but constitute distinct sources of evidence for the independence of ornaments.

#### 7.2 The first $(n-1)$ clicks of ornamented codas are more similar to neighbouring codas' rhythm type than any $n$-click rhythm type

Codas are grouped into rhythm types based on the number of clicks in the coda and the relative spacing of the clicks in their standardised form. However, in the case of ornamented codas, the ornament appears to be independent of the rest of the clicks in the coda, and the remaining clicks together form a coda with a rhythm type matching to the neighbouring codas by the same whale. Consider a standardised ornamented coda:

$$\mathbf{c}_o = [t_2 - t_1, t_3 - t_1, t_4 - t_1, t_5 - t_1, t_6 - t_1]/(t_6 - t_1)\ . \tag{4}$$

The coda's standardised representation with the ornament-removed is:

$$\mathbf{c}_{-o} = [t_2 - t_1, t_3 - t_1, t_4 - t_1, t_5 - t_1]/(t_5 - t_1)\ . \tag{5}$$

Let `clusterCentre` denote the function that returns the cluster centroid of a standardised coda type, and let `neighbour` denote the function that returns a standardised neighbouring coda made by the same whale. We compare the distributions (as seen in Fig. 15):

1. Quantity 1 (in orange): $\|\texttt{clusterCentre}(\mathbf{c}_o) - \mathbf{c}_o\|_2^2$

2. Quantity 2 (in blue): $\|\texttt{clusterCentre}(\texttt{neighbour}(\mathbf{c}_o)) - \mathbf{c}_{-o}\|_2^2$

The average value of quantity 2 (0.0034s$^2$) is significantly less than the average value of quantity 1 ($0.0053s^2$) (test: permutation test (one-sided), $p = 0.0022$, $n = 178$, performed by randomly resampling labels [distance to cluster vs distance to neighbor's cluster]; Figure 15). Therefore, the internal distribution of clicks in an ornamented coda is more like its neighbouring codas than the coda cluster it is originally assigned to.

#### 7.3 Ornamented codas have a higher variance in the difference of durations between adjacent codas by a given whale

Generally, in the course of an exchange, a whale fixes the rhythm that it makes, picks a tempo type to start with, and only gradually varies the duration of the coda. However, ornamented and non-ornamented codas exhibit significant differences in the distribution of their duration differences with neighboring codas (test: Kolmogorov-Smirnov test (two-sample), $D(206, 206) = 0.25$, $p = 5e^{-11}$, 95% CI= $[0.15, 0.35]$, $n = 206$).

#### 7.4 ICI values of ornamented codas are atypical

As described in Section 2.1 of the main paper, the difference in the two final ICIs, normalized by the duration of the penultimate ICI, exhibits a significantly different distribution in ornamented and non-ornamented codas (test: Kolmogorov-Smirnov test (two-sample), $D(178, 3666) = 0.28$, $p = 2e^{-14}$, 95% CI= $[0.17, 0.39]$, $n = (178, 3666)$; main paper Fig. 2(D)).

#### 7.5 Ornaments are not imitated

In overlapping codas where at least one of the codas is ornamented, the number of clicks the ornamented coda is only matched (5%) of the time, compared to (71%) of the time for non-ornamented overlapping codas. The average difference in duration for overlapping codas of which at least one is ornamented (0.210s) is also significantly larger than the difference in duration of overlapping non-ornamented codas (0.099s) (test: permutation test (one-sided), p=0.0001, n=(62, 848), performed by randomly resampling labels [ornamented vs. unornamented] without replacement).

#### 7.6 Ornaments are non-uniformly distributed across call sequences

If the ornamentation were an uncontrolled feature in sperm whale calls, it would be equally likely to occur anywhere in the sequence and therefore, uniformly distributed as a function of time in the call sequence. However, ornaments are more likely to be present at the beginnings and ends of call sequences. For this analysis, we consider call sequences with greater than two codas in the sequence. Fully more than half of ornaments occur at the beginning or end of a call sequence: (29.08%) in the first coda and (26.24%) in the last coda. Coda type counts by location are given in Table 1.

| | Start of sequence | Elsewhere |
|---|---|---|
| Ornamented | 41 (29%) | 100 (61%) |
| Non-ornamented | 594 (17%) | 2893 (83%) |

| | End of sequence | Elsewhere |
|---|---|---|
| Ornamented | 37 (26%) | 104 (74%) |
| Non-ornamented | 598 (17%) | 2889 (83%) |

| | Before chorusing change | Elsewhere |
|---|---|---|
| Ornamented | 82 (66%) | 42 (34%) |
| Non-ornamented | 1465 (50%) | 1434 (50%) |

Table 1: Frequency of ornamented and non-ornamented codas at the start and end of coda sequences (top and middle respectively), and frequency before changes in vocalization style (bottom)

From this table, it can be seen that ornamented codas are more likely to occur at the start of a whale's calls compared to non-ornamented codas (test: Fisher's exact test (two-sided), odds ratio: 2.00, $p$ =0.0006). Ornamented codas are also more likely to occur at the end of a whale's calls as opposed to non-ornamented codas (test: Fisher's exact test (two-sided), odds ratio: 2.07, $p$ =0.0004).

#### 7.7 Ornamented codas are followed by a change in responding whale vocalizations

We define a 'change in chorusing behavior' as one of three events: a following whale begins chorusing with a leading whale, pauses chorusing, or ceases vocalizing for the remainder of the exchange. Counts by location relative to chorusing change events are also shown in Table 1. Compared to unornamented codas, ornamented codas from the *leading* whale are disproportionately succeeded by a change in chorusing behavior from the *following* whale (test: Fisher's exact test (two-sided), odds ratio: 1.91, $p = 0.0007$).

| **Traits in Human Language** | **Sperm Whales** |
|---|---|
| Vocal auditory channel | Yes. (*10*) |
| Broadcast transmission, and directional reception | Yes. (*10, 11*) |
| Rapid fading (Transitoriness) | Yes. |
| Interchangeability | Yes. |
| Total Feedback | Yes. |
| Specialization | Yes. (*11*) |
| Semanticity | Maybe. (*11, 12*) |
| Arbitrariness | Yes. (*1, 10, 11, 13, 14*) |
| Discreteness | Yes. (*9, 10*) |
| Displacement | ? |
| Productivity | ? |
| Traditional Transmission | Yes. (*1, 13, 14*) |
| Duality of Patterning | Maybe. [This paper] |

Table 2: Characteristics of Language: Hockett's Design Features of Natural Language. A detailed description of each of the listed features can be found in (*15*).

### 8 Information Capacity

#### 8.1 Fixed coda inventory

Previously, the communication system of the Eastern Caribbean sperm whales was hypothesized to consist of 21 unique coda types. If at a given time a single coda is produced, such a communication system could communicate at most $\lceil \log_2 21 \rceil = \lceil 4.39 \rceil = 5$ bits per coda.

#### 8.2 Combinatorial coda inventory

The proposed system hypothesizes that codas are constructed by sampling from 18 rhythm types, the presence or absence of an ornament (2 possible values), an increasing, decreasing, or unchanging rubato (3 possible values), and one of 5 tempo types. The resulting set of codas that can be generated from such a system can contain a total of $18 \times 5 \times 2 \times 3 = 540$ symbols, capable of communicating $\lceil \log_2 576 \rceil = \lceil 9.08 \rceil = 10$ bits per coda. Therefore, the set of new features and a combinatorial coding system allows encoding a significantly greater number of messages.

However, not all possible codas are frequently realized in practice. 20 of the rhythm and tempo type combinations occur with both rubato and ornamentation, 6 with only rubato, 1 with only ornamentation, and 16 without ornamentation or rubato. Here we ignore infrequent occurrences ($\leq 1$ time in the DSWP dataset) of features across the different types. The resulting set of frequently generated feature combinations comprises 156 different codas, capable of com-

| Feature | Human Language |
|---|---|
| Phonetic features | Tonality, pitch, rate, nasality, etc |
| Phonemes | /b/, /d/, /f/, /g/, etc |
| Morphemes | *eat, date, weak, -ness, -ly, -ism, etc* |
| Words | *eats, dated, weakness, happily, etc* |
| Sentences | *Lily ate some fruit.* |

Table 3: The linguistic hierarchy in humans and instances of the features in English.

municating $\lceil \log_2 156 \rceil = \lceil 7.28 \rceil = 8$ bits per coda.

A computation based on the size of the message space alone is an upper bound on the true information rate. This is because, in practice, calls made at consecutive time steps are not independent. Consecutive calls made by a given whale are more likely to be of the same rhythm type (Section 4) and overlapping calls are likely to have the same overall duration (Section 6). Therefore, computing the information transmission capacity of the communication system requires accounting for dependencies between calls over longer time horizons, an important avenue for future work.

### 9 Parallels to Human Language

While an animal's communication system does not have to be human-like to exhibit non-trivial structure, human language is considered the epitome of structured communication systems due to a combination of regularity and expressivity. A comparison between the human language and sperm whale communication allows for benchmarking aspects of the (incompletely characterized) whale communication system against well-studied aspects of human communication. Inspired by the study by (*15*), Table 2 highlights the design features of natural language and what has been understood of them in the sperm whale communication system. Table 3 and Table 4 highlight features of the linguistic hierarchy in human language and their possible corresponding analogs in sperm whale communication thus far. Aspects of this characterization are provisional, and further study and analysis of the sperm whales' communication system will allow for continued in-depth characterization.

### 10 Additional Discussion of Statistical Tests

Comparisons of coda durations (either with adjacent codas, when studying rubato, or with overlapping codas, when studying all features in the context of chorusing behavior) to avoid making

| **Linguistic Features** | **Analogues** | **Examples** |
|---|---|---|
| Phonetic features<br>Phonemes | Sub-coda features | Rhythm, Tempo, Rubato, Ornamentation |
| Morphemes / Words | Codas | 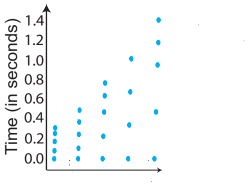 |
| Sentences | Coda Sequences | 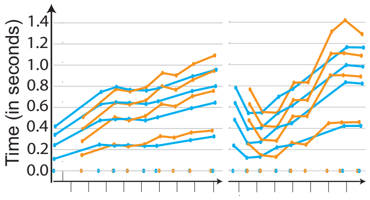 |

Table 4: Analogues of the features of the linguistic hierarchy in sperm whale communication.

distributional assumptions about durations of codas and their absolute differences, some of which have non-normal distributions. All permutation tests computed over 10,000 random resamplings of the data without replacement. Evaluation of rubato additionally uses Spearman rank-correlation tests to measure longer range trends across coda triplets (again based on initial observations that these trends appeared to be non-linear). Comparisons of coda-internal structure (e.g. durations of penultimate ICIs in ornamented codas) use Kolmogorov-Sminrnov tests, as we are interested only in distributional differences rather than orderings of mean durations. Finally, measurements of changes in vocalization behavior following rubato use Fisher's exact test (another permutation test) to compare proportions of these changes in different vocal contexts.

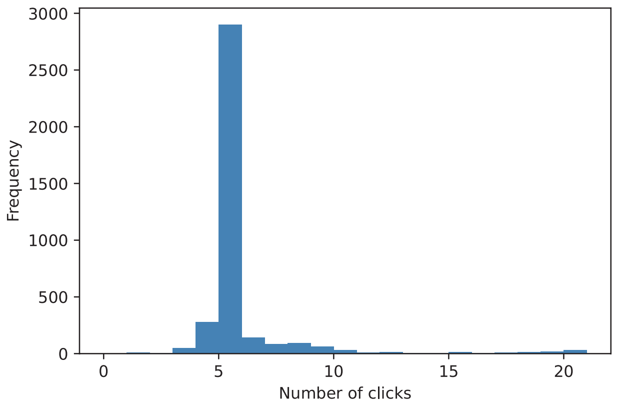


Figure 1: Frequency distribution of codas based on the total number of clicks in the coda for the Dominica Sperm Whale Project Dataset. Sperm whales of the Eastern Caribbean Clan-1 predominantly produce 5-click codas.

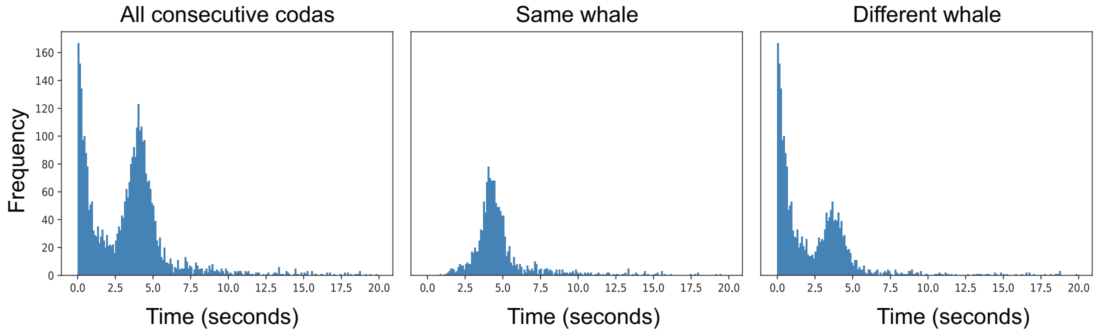


Figure 2: Response time: In the EC1 clan, consecutive codas are produced such that they either overlap with one another (response time $\approx 0$) or after a time period of four seconds. *(Left)* The frequency distribution of the response time between consecutive codas. *(Center)* The frequency distribution of the response time if the consecutive codas were made by the same speaker, denoting the four-second isochronous periodicity of the coda production of an individual whale; and *(Right)* The frequency distribution of response time when consecutive codas are produced by different whales. The peak at $t = 0$ corresponds to overlapping codas, and the peak at $t \approx 4$ corresponds to codas produced during turn-taking.

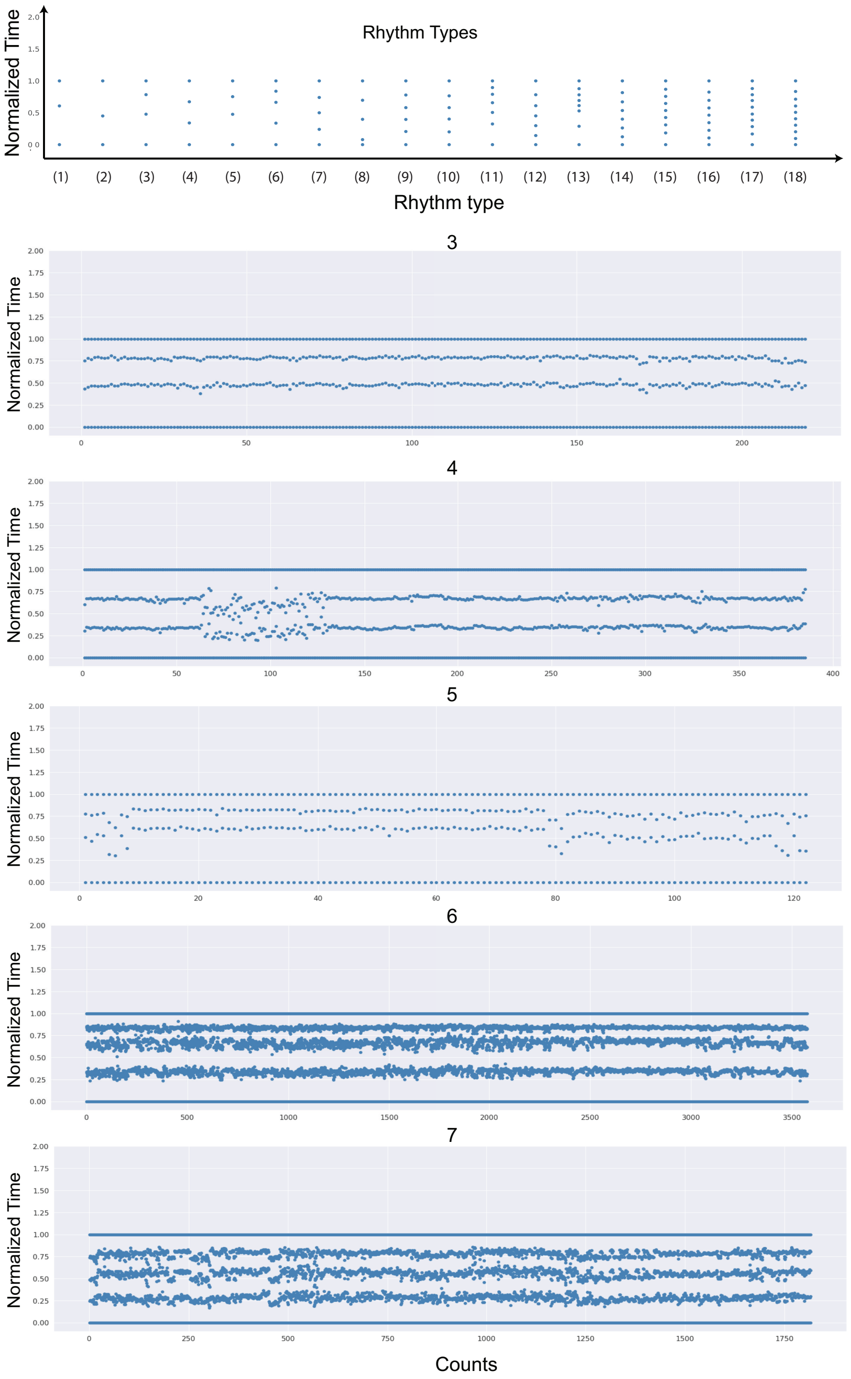


Figure 3: *(Top)* The set of coda rhythm types denoted by the mean standardized coda in each of the clusters. *(Bottom)* The corresponding set of codas of rhythm types 3 to 7.

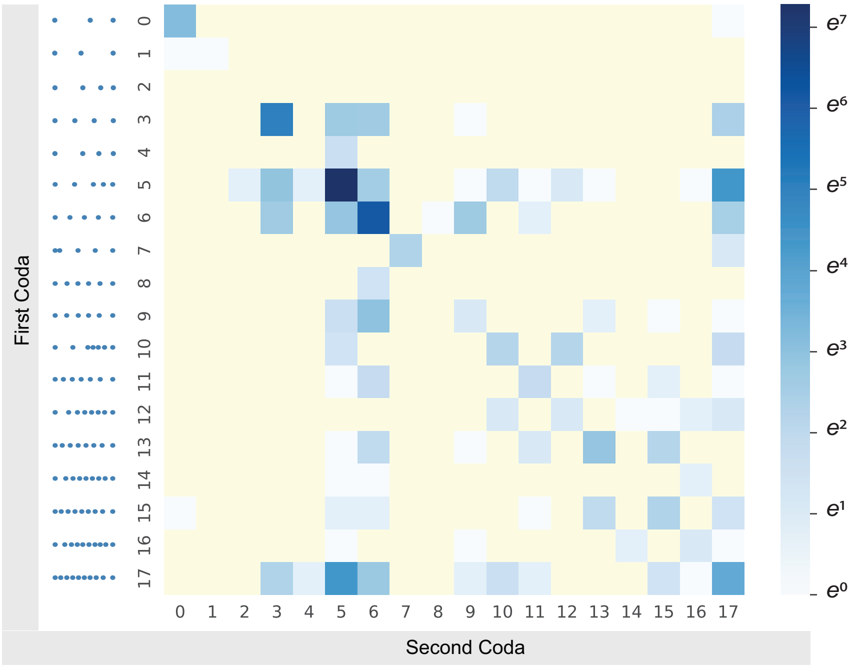


Figure 4: Frequencies of transitions between different rhythm types.

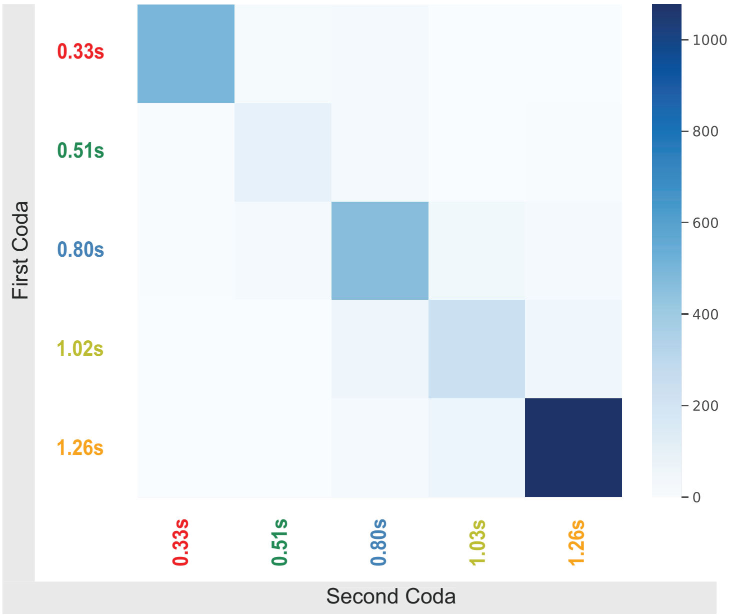


Figure 5: Frequencies of transitions between different tempo types. The matrix is diagonally dominant.

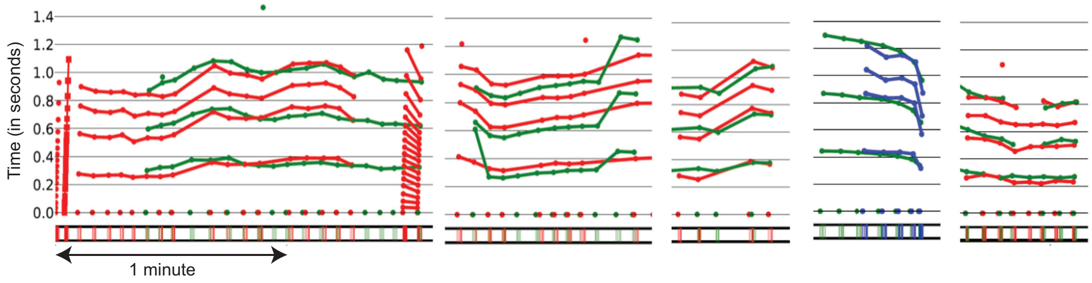


Figure 6: A set of exchange plots showing choruses between pairs of sperm whales. Here, whales exchange overlapping codas of different rhythm types while matching duration. This demonstrates independent control of rhythm and duration during coda production.

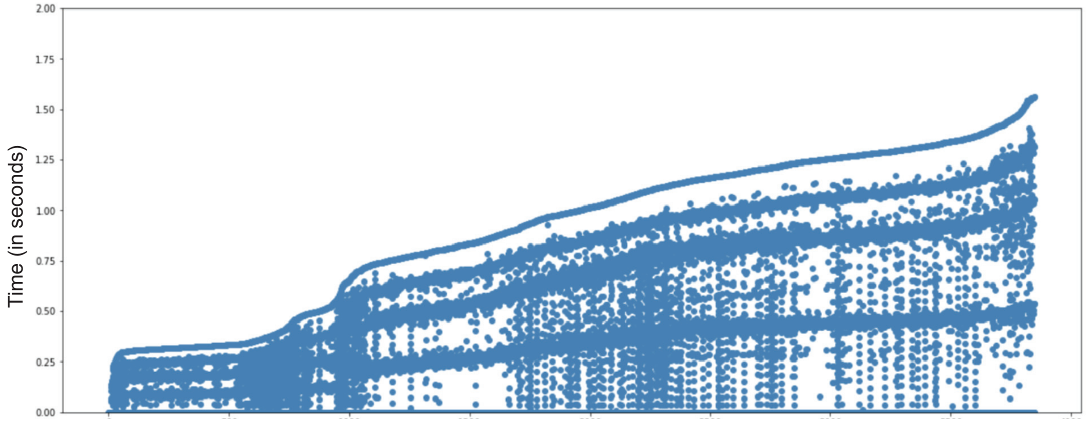


Figure 7: All DSWP codas visualized in increasing order of overall duration. The empirical distribution of the duration cluster around a set of small modes as seen in Fig. 2(B) of the main manuscript.

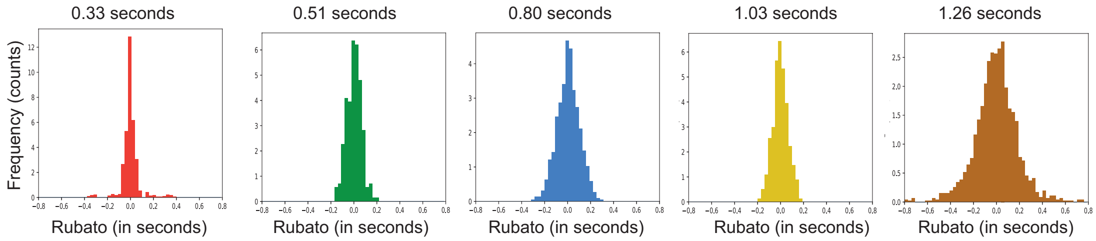


Figure 8: Distribution of rubato between adjacent codas for each of the 5 tempo types. The rubato for each of the tempo types is unimodally distributed around a mean value of zero.

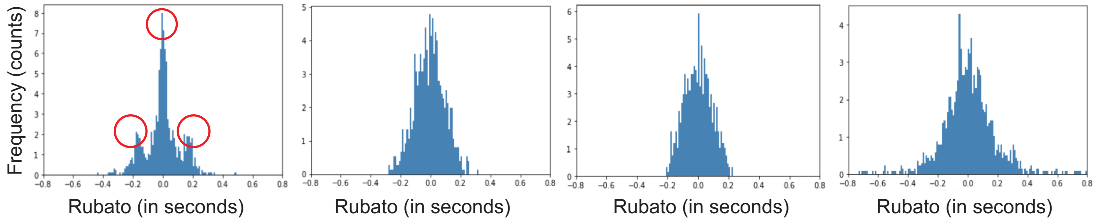


Figure 9: A reduction in the number of tempo clusters to four results in a multi-modal distribution of rubato.

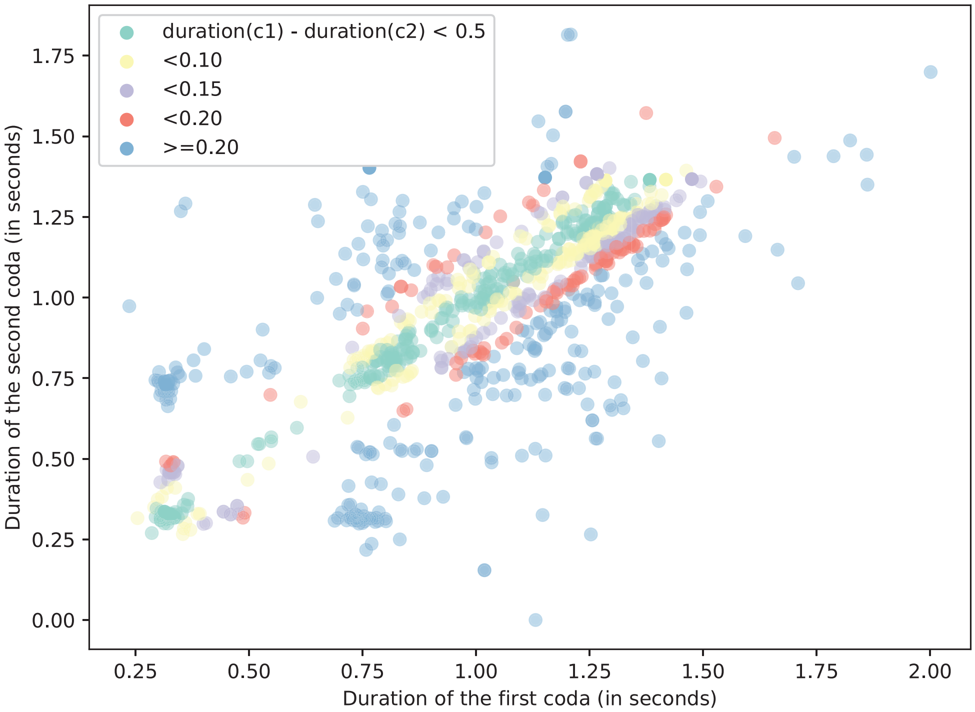


Figure 10: Overall durations of overlapping codas in the dataset. Overlapping codas are imitated with varying degrees of precision. The breakdown of the percentage of overlapping codas in each category and its corresponding precision range are: green: (25%) within 0.05 seconds; yellow: (45%) within 0.1 seconds; purple: (61%) within 0.15 seconds; red: (71%) within 0.2 seconds; blue: remaining set of overlapping codas (100%).

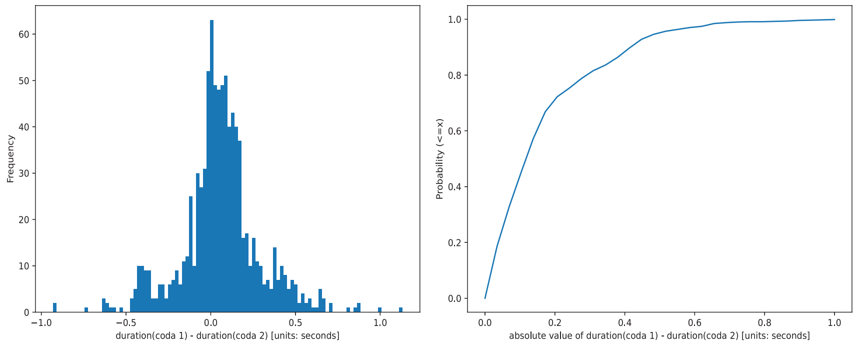


Figure 11: (*Left*) The difference in durations of overlapping codas is distributed tightly around zero. This indicates that the durations of codas are predominantly closely matched. (*Right*) The cumulative distribution function of the difference in durations of overlapping codas. The CDF indicates that about (70%) of the overlapping codas are matched with a precision of about $(1/5)^{th}$ of sec.

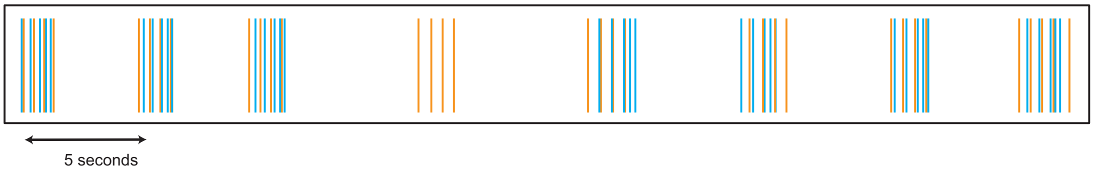


Figure 12: Overlapping codas made by a pair of whales in a chorus. In this exchange, the first click of the second whale in each pair of overlapping coda is made between the first two clicks of the coda of the leading whale.

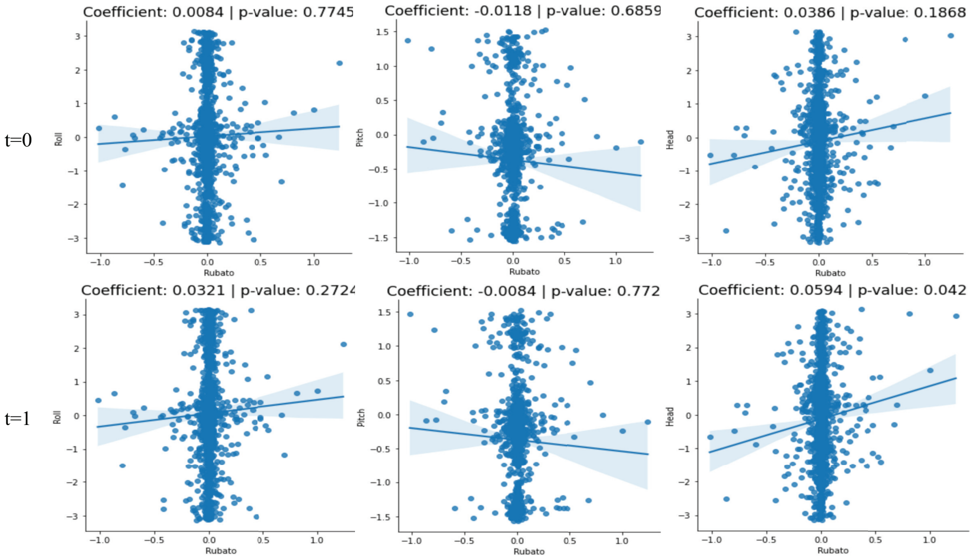


(i) Rubato versus Gyroscope readings

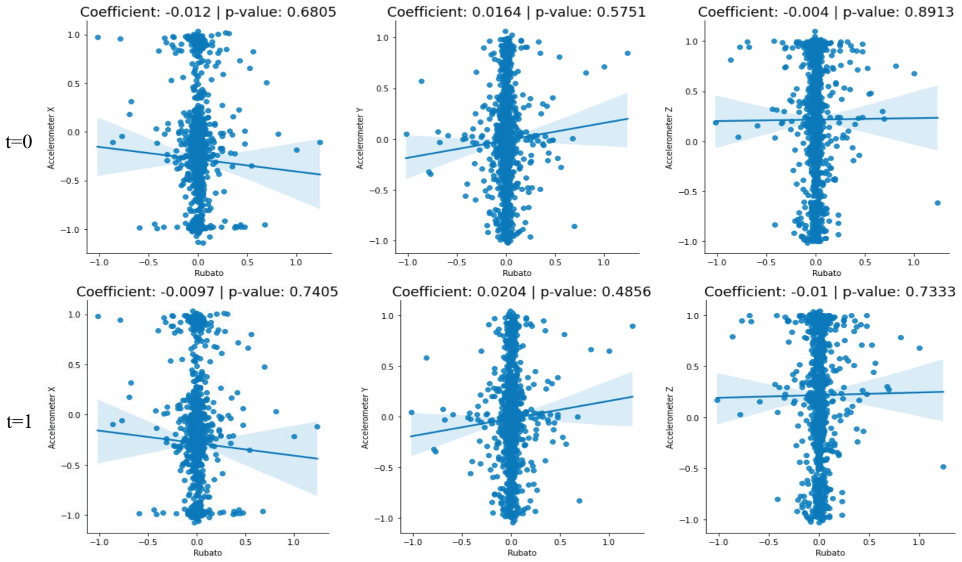


(i) Rubato versus Accelerometer readings

Figure 13: No significant correlation was found between gyroscope readings and rubato or between the accelerometer readings and rubato. The correlation between the quantities was checked from orientation and acceleration measurements made zero seconds apart to motion readings up to 10 seconds before (test: Spearman's rank order correlation test (two-sided), $n = 1172$).

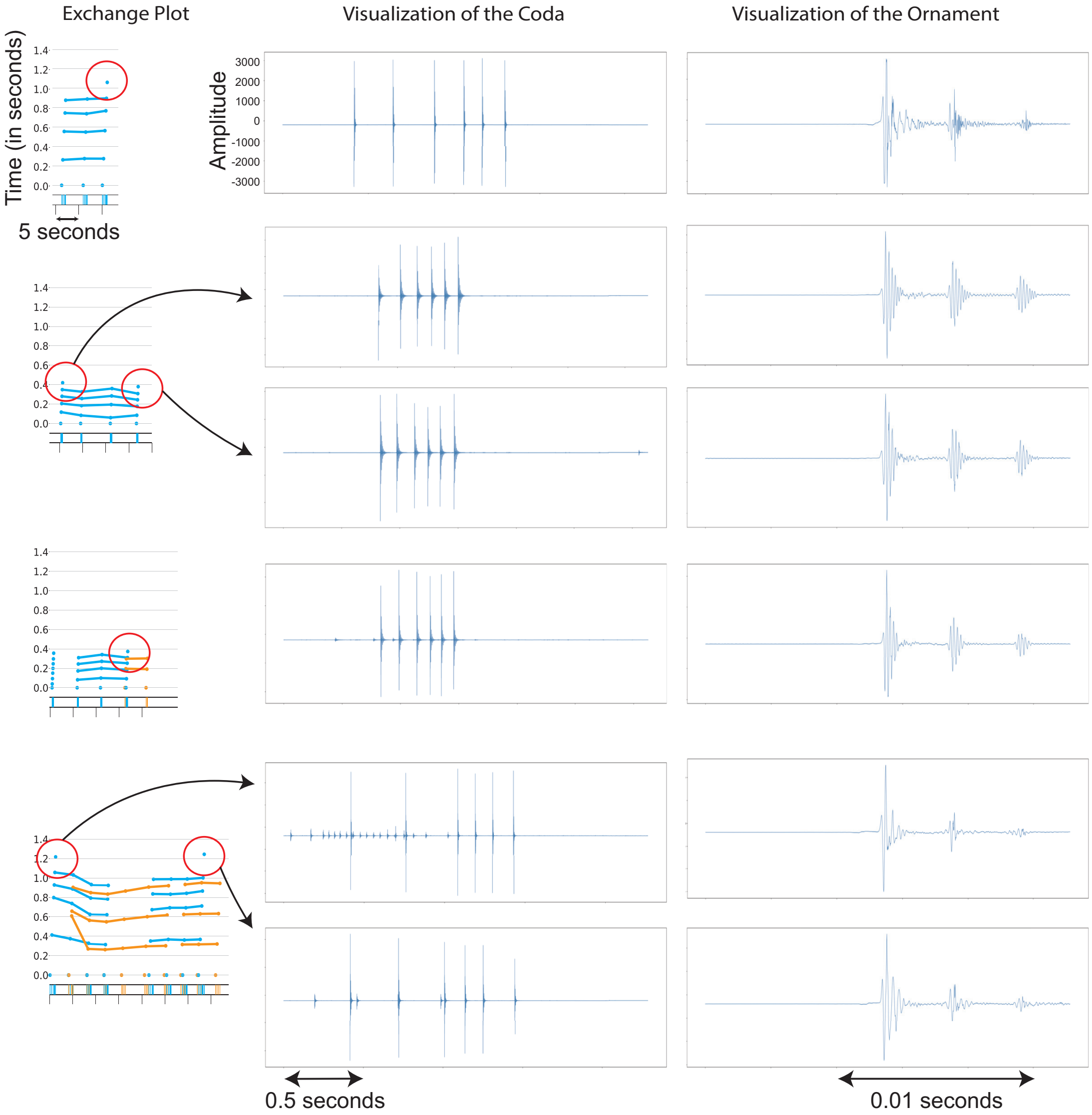


Figure 14: *(Left)* A snippet of an exchange plot where one of the whales produces an ornament. *(Center)* The waveform of the ornamented coda. *(Right)* The waveform of the ornament. The ornament is structurally similar to the remainder of the clicks in the coda.

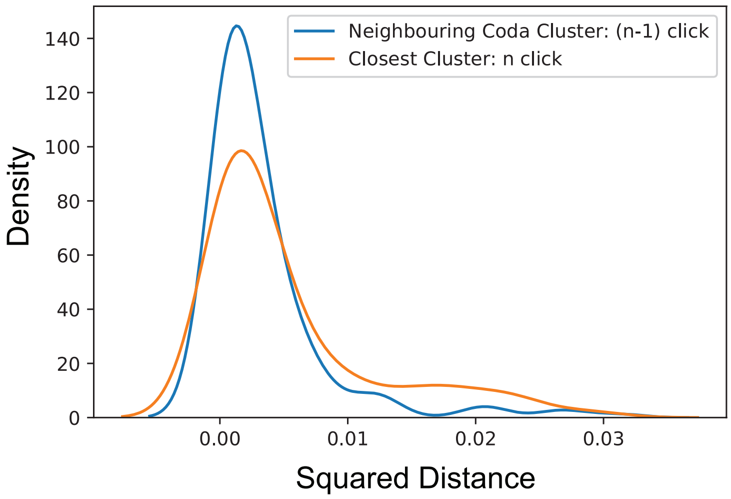


Figure 15: Distribution of distances of (orange) standardised ornamented codas with the standardised cluster centers of their rhythm types, and (blue) standardised ornamented codas with the ornament removed with the cluster centers of the neighbouring codas. Rhythm types of the first (n-1) clicks of ornamented codas are closer to those of the neighbouring codas than the ornamented coda's rhythm type is to its assigned cluster.

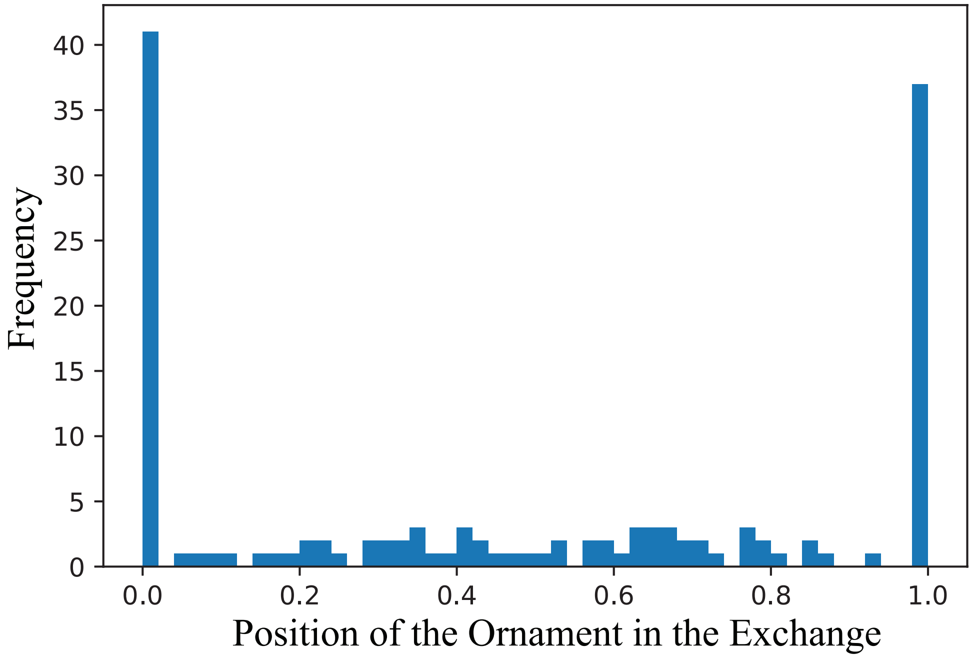


Figure 16: Frequency distribution of ornamented codas in the span of consecutive call sequences made by an individual whale. Calls were defined to be consecutive if they occurred no later than 8 seconds apart (which is twice the average inter-coda time as described in Figure 2 of the Supplementary Material).
